# A transient burst of mutations occurs during the normal development of yeast colonies

**DOI:** 10.1101/2023.12.11.571082

**Authors:** Nicolas Agier, Nina Vittorelli, Frédéric Chaux, Alexandre Gillet-Markowska, Samuel O’Donnell, Gilles Fischer, Stéphane Delmas

## Abstract

Characterizing the pace of mutation accumulation is crucial for understanding how populations adapt to their environment and for unraveling the intricate dynamics between gradual processes and more sudden burst-like events occurring during cancer development. We engineered the genome of *Saccharomyces cerevisiae* to measure the rates of single and double mutations, including point mutations, segmental duplications and reciprocal translocations. We found that during the development of wild-type yeast colonies, double mutations occur at rates that are up to 17-fold higher than those expected on the basis of single mutation rates. We found that this excess of double mutations is partially dependent on the *ELG1/ATAD5* clamp unloader. Additionally, the double mutants retain wild-type mutation rates, suggesting that they originated from genetically wild-type cells that transiently expressed a mutator phenotype. Numerical simulations based on the experimentally measured mutation rates, confirmed that the excess of double mutations can be accounted for by subpopulations of transient mutators within the colony. These subpopulations would be limited to less than a few thousand cells and temporarily adopt mutation rates multiplied by hundreds or thousands for less than five generations. We found that the majority of double mutations would accumulate sequentially in different cell cycles. The simultaneous acquisition of both mutations during the same cell cycle would be rare and possibly associated with systemic genomic instability. In conclusion, our results suggest that transient hypermutators play a major role in genomic instability and contribute significantly to the mutational load naturally accumulating during the growth of isogenic cell populations.

**Significance statement:** Understanding the pace at which mutations accumulate is of paramount importance in the field of genome dynamics and evolution. In our study, we unveiled a surprising burst of mutations within growing yeast colonies, occurring independently of external stressors. This discovery indicates that, during short intervals, a small subset of cells within the colonies undergoes a mutational overdrive. Notably, these mutator cells do not represent genetically stable mutators with mutations in genes associated with genome stability. Instead, they stem from a strong mutator phenotype that was transiently expressed in genetically wild-type cells. This phenomenon, previously underestimated or even overlooked, holds significant importance and may have far-reaching implications, particularly in the context of cancer development.

## Introduction

Mutations arise through errors during the transmission of the genetic material over the generations, leading to permanent changes in the DNA sequence. Unfaithful repair of DNA replication errors and DNA damage induced by endogenous or exogenous stresses such as reactive oxygen species or UV, respectively, are among their main causes (1). The regime and tempo of accumulation of mutations in a genome has a direct impact on the capacity of a population to adapt and to evolve. It has long been assumed that the mutation rate remains constant and is kept low but not null in a population due to a balance between various factors such as the prevalence of deleterious mutations over beneficial ones, the energetic cost of protein fidelity or the drift barrier at which even lower mutation rates are not more beneficial (2–4).

However, over the last decade, the mutation accumulation model has departed from the classic regime of gradual accumulation of independent mutations over time towards a more complex model in which periods of mutation explosion occur between periods of gradual accumulation (5). These bursts of mutation were characterized by the presence of mutants carrying more mutations than expected based on the mutation rate (6). These mutations were either clustered in the genome or dispersed over long distances along chromosomes (6). It was shown that, in changing environments, the cells that acquire a higher mutation rate, referred to as mutators, can be favored through natural selection because of their increased likelihood of generating adaptive mutations (3, 4, 7–10). Mutators are said to be genetic when they result from mutations in genes involved in genome maintenance. Genetic mutators are of great importance, from acquisition of antibiotics resistance in bacteria, to drivers mutations in tumorigenesis (1, 11). However, after adaptation, genetic mutators progressively lose fitness due to the increased accumulation of deleterious mutations and eventually become extinct unless they revert to lower mutation rates by obtention of rare reversion or suppressor mutations (12, 13). By opposition to the genetic mutators that stably exert their effect over generations, another type of mutator cells called phenotypic or transient mutators was described. A semi-quantitative analysis of the contribution of transient phenotypic mutators to single and double mutations in *Escherichia coli* suggested that most double mutations would result from transient mutators (Nino et al. Genetics, 1991). In bacteria and yeast, it has been shown that a transient mutator phenotype can originate from cell to cell heterogeneity in response to intrinsic or extrinsic stresses (14–16). Fluctuation in the quantity of low copy number proteins involved in DNA replication and repair due to unequal segregation at division, or errors during the transcription or translation process that would cause loss of function or toxic gain of function mutant proteins can cause a temporary mutator phenotype until the protein balance is restored (6, 17, 18). For instance, in a clonal population of diploid *Saccharomyces cerevisiae* cells, a detailed assessment of multiple instances of loss of heterozygosity events and chromosome copy number alterations revealed the existence of subpopulations of cells capable of experiencing transient systemic genomic instability (SGI) (19, 20). Thus, bursts of mutations could result from the transient coexistence of heterogeneous mutation rates inside a clonal cell population (7, 9). However, many questions concerning the size of the mutator subpopulations, the fold increase of the mutation rate (mutational strength), the duration of the mutator episodes as well as their underlying molecular causes remain unanswered.

In this study, we discovered the existence of transient mutator subpopulations that influence the regime of accumulation of mutations during the normal development of wild-type yeast colonies grown in complete media and in absence of exogenous stress. Our experimental results, complemented with numerical simulations, suggest the presence of mutator subpopulations of small size and high strength that strongly increase the rates of both sequentially and simultaneously acquired double mutations. Our results suggest that the transient hypermutator phenotype does not arise from DNA replication defects but is partly dependent on the Elg1/ATAD5 clamp unloader and so potentially related to the recombination salvage pathway that allow bypass of lesions at single-stranded gap (21).

## Material and Methods

### Strains, growth media and reagents used in this study

All strains used in this study were derived from the haploid BY4741 (*MAT a, his3Δ1 leu2Δ0 met15Δ0 ura3Δ0*) (22) and presented in **supplementary table 1**. *RAD52*, *RAD27* and *ELG1* gene deletions were constructed by transformation of a PCR product containing the *KanMX4* selectable marker flanked by 50 pb of the upstream and downstream sequences of the targeted gene (23). Gene deletions were verified by growth on selective media, amplification of the insertion region by PCR and validation by sanger sequencing. Genetic reporter systems to measure the rate of segmental duplication (*D*) described in (24), reciprocal translocation (*T*) described in (25) and the resistance to the canavanine (*C*) are presented in **Figure 1A**. Cells were grown on empirical YPD (broth or agar) (Sigma-Aldrich) or synthetic media (glucose 20 g/L (Sigma-Aldrich), Yeast Nitrogen Based 6 g/L (Sigma-Aldrich), agar 20 g/L (Sigma-Aldrich), and complemented with specific amino acid mixes according to the required selection. These mixes were either the “drop-out’’ (Sigma-Aldrich) or “CSM” (MP Biotech). When required uracil, leucine, canavanine or hydroxyurea (all from Sigma-Aldrich) were added at 76 mg/L, 380 mg/L, 60 mg/L or 100 mM respectively.

**Figure 1:**
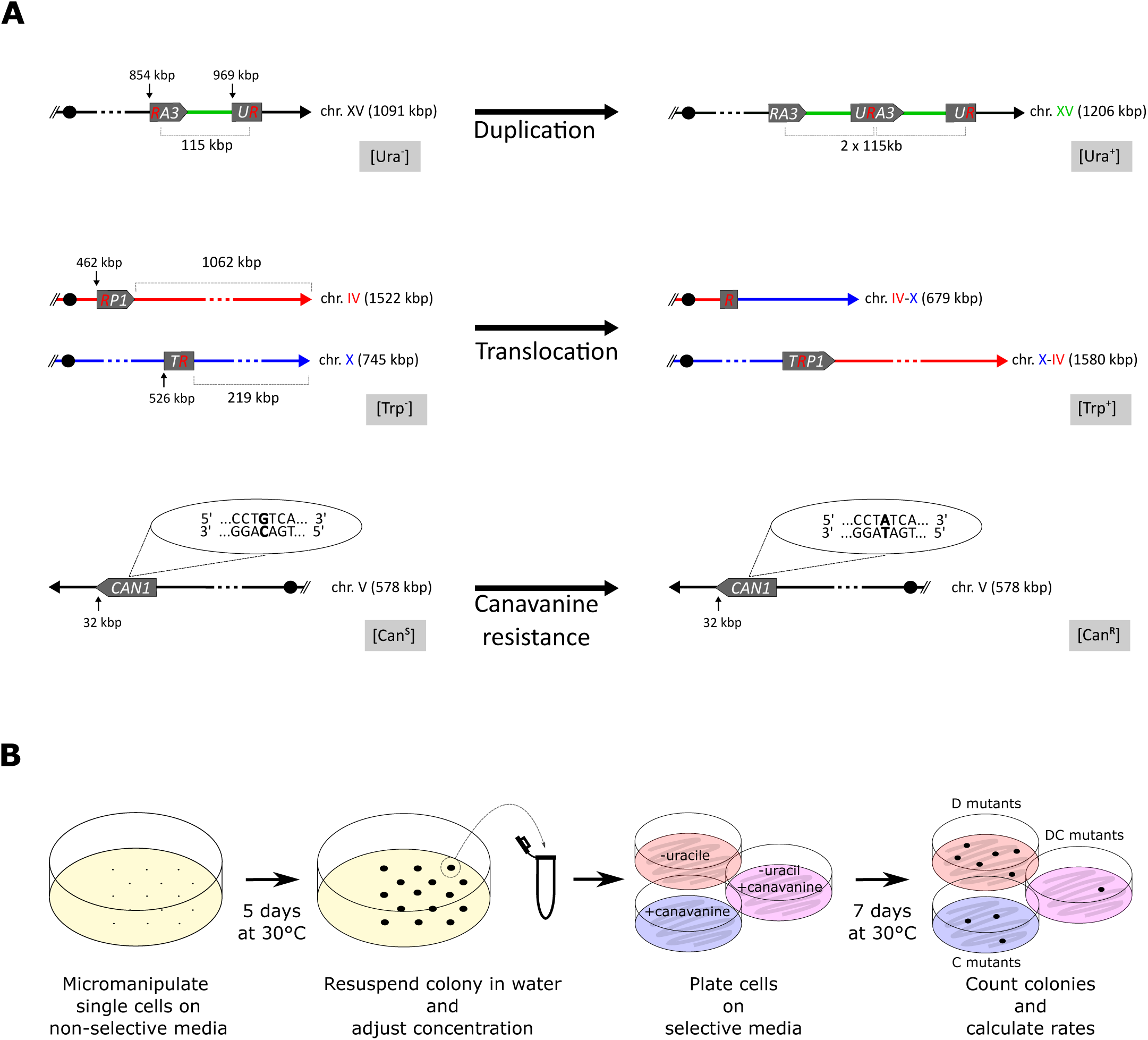
Reporter systems and experimental protocol. **A)** The <u>D</u>uplication system (*D*, top) is composed of two split alleles, “*RA3*” and “*UR*”, deleted from the 5’ and 3’ part of the *URA3* gene, respectively. These alleles are respectively located at coordinates 854 kb and 969 kb on chromosome XV. The two alleles share an overlapping region of 450 bp promoting, by homologous recombination, the formation of a 115 kb segmental duplication and the restoration of a functional *URA3* gene at its junction. The duplication leads to an increase in size of chromosome XV to 1206 kb and a phenotypic switch from uracil auxotrophy [Ura^-^] to prototrophy [Ura^+^] allowing mutant cells to be selected on a synthetic medium lacking uracil. The <u>T</u>ranslocation system (*T*, middle) is composed of two split alleles, “*RP1*” and “*TR*”, deleted from the 5’ and 3’ part of the *TRP1* gene and located at coordinates 462 kb and 526 kb on chromosomes IV and X, respectively. These two alleles share an overlapping region of 450 bp promoting, by homologous recombination, the formation of a reciprocal translocation and at its junction a functional *TRP1* gene. The reciprocal translocation leads to the formation of two new chromosomes IV-X and X-IV and a phenotypic switch from tryptophan auxotrophy [Trp^-^] to prototrophy [Trp^+^] allowing the selection of mutant cells on a synthetic medium lacking tryptophan. The <u>C</u>anavanine system (*C*, bottom) relies on the counter-selectable marker *CAN1* gene located at coordinates 32 kb on chromosome V. Can1 is a native permease allowing the uptake of the lethal arginine homolog, canavanine (ref?). Mutation in *CAN1* (symbolized here by a G to A substitution) prevent canavanine uptake and thus this loss of function leads to a phenotypic switch of the cells from canavanine sensitivity [Can^S^] to resistance [Can^R^] and allows mutant cells to be selected on a synthetic medium lacking arginine and supplemented with canavanine. **B)** Cells are deposited at equal distance from each other using a micromanipulator on a non-selective complete synthetic medium and incubated for 5 days at 30°C. The colonies are recovered in 200 µL of sterile water, diluted if required and plated on three selective synthetic media, (i) lacking uracil, (ii) with addition of canavanine and (iii) lacking uracil and with the addition of canavanine to select for *D, C* or *DC* mutants, respectively. The selection for *T* mutants on a medium lacking tryptophan is not represented here. The selective plates are incubated for 7 days at 30°C before counting the mutant colonies.

### Measurement of mutation rates on structured and liquid medium

Fluctuation assays were performed to measure the mutation rates on structured or liquid media. On structured medium, strains were destocked from −80°C on YPD agar and allowed to grow at 30°C. After 24 hours, using a micromanipulator, 15 cells per plate were individually deposited, at equal distance from each other, on a glucose supplemented synthetic complete agar medium and the plates incubated at 30°C for 5 days (**Figure 1B**). Colonies, with the exception of petite mutant colonies which we excluded, were individually recovered in 200 µL of sterile distilled water, diluted if required, and plated on appropriate selective synthetic medium to select for single or double mutations. In each experiment, to estimate the number of cells in a colony, we measured the optical density at 600 nm (OD_600_) for 8 random colonies. The selective plates were incubated for 7 days before colonies were counted, and mutation rate calculated using r-salvador (26). To compare two mutation rates, we applied the likelihood ratio test method using r-salvador (26). To measure the mutation rates in liquid medium, strains were destocked from −80°C on YPD agar and allowed to grow at 30°C. After 24 hours a suspension of cells is prepared on complete synthetic medium, the concentration of cells determined by measuring the OD_600_ and cultures inoculated with an average of 10 cells. Cultures were incubated at 30°C with 150 rpm agitation for 3 days. Then the same protocol as in structured medium is applied to select for mutant colonies. All mutation rates are presented in **supplementary table 2**.

#### Measurement of the generation time

Strains were destocked from −80°C on YPD agar and allowed to grow at 30°C. After 48 hours, growth curves were performed by inoculating 10^5^ cells in 200 μL in YPD broth on a 96 wells plate. The plate is then incubated at 28°C without agitation in a TECAN sunrise spectrophotometer. Measurements of the OD_600_ were taken each 10 minutes. The generation time was established using the R package GETITshiny 3.4 available on GitHub (DOI. 10.5281/zenodo.10227629).

### Pulse Field gel Electrophoresis (PFGE)

Whole yeast chromosome agarose plugs were prepared according to (27) and sealed in a 1% Seakem GTC agarose and 0.5x TBE gel. PFGE was performed with a CHEF-DRIII (BioRad) system with the following program: 6 V/cm for 10 hours with a switching time of 60 seconds followed by 6 V/cm for 17h with switching time of 90 seconds. The included angle was 120° for the whole duration of the run. Agarose gels were stained with ethidium bromide for 20 minutes in TBE 0.5X.

### PCR and sequencing CAN1 gene

To determine the mutation spectrum the *CAN1* gene was amplified and sequenced by the Sanger method at Eurofins using primers “CAN1extR2” and “CAN1ext/intF1” (28).

### Long-read sequencing, de novo assembly and genome analyses

Genomic DNA was extracted using QIAGEN genomic TIP 100/G, following the manufacturer’s protocol. Size selection was done using Circulomics SRE kit. DNA from 10 strains was then prepared using the ONT LSK109 kit coupled with the EXP–NBD104 barcoding kit and sequenced on a single R9.1 flowcell using a MK1C Minion sequencer. Standard High Accuracy base calling was performed on the Mk1C with the 22.05.8 version of minknow, using Guppy (V6.1.5, model=dna_r9.4.1_450bps_hac.cfg). For data processing, barcodes were removed from sequences using porechop. Then sequences were downsampled using Filtlong to obtain a coverage of 40X enriched for reads with high quality and longer length. Genome assembly was performed using 3 different assemblers : Canu (v2.1), SMARTdenovo (https://github.com/ruanjue/smartdenovo/ v1.12) and NextDenovo (https://github.com/Nextomics/NextDenovo). Contigs were polished using 1 to 3 rounds of Racon (v1.4.21) and 2 rounds of Medaka (https://github.com/nanoporetech/medaka v1.2.3). Finally reference Scaffolding was performed with Ragout (v2.3). SVs were called using MUM&Co (v2.4.2) (29), on all 3 assemblies. We validated SVs found in at least 2 assemblies. We also checked that we detected the expected deletions corresponding to the auxotrophic markers in the BY4741 background and the insertions of resistance cassettes when applicable. For chromosomal rearrangements, manual scaffolding was performed and validated by karyotyping with PFGE. SNPs were called using medaka_variant. We only considered haploid SNPs with a minimum quality score of 45 which corresponded to the lower quality score of the selected *CAN1* mutations that were validated by Sanger sequencing. The fastq files are available on the ENA website under the accession number PRJEB70654/ERA27514923.

### A stochastic model to estimate the double mutation rate

Both null and refined versions of the model are available in the SIMPLE_GUI_Neutral.R package (https://github.com/Nico-LCQB/SIMPLE_GUI, DOI 10.5281/zenodo.8325449). For both models, the number of generations and the values of the two single mutation rates are chosen by the user. For the refined model, the size of the mutator subpopulation, the fold-increase of the mutation rates and the duration of the mutator episode (in generations) are also defined by the user. The output provides the following information: (i) the number of each type of single and double mutations that occurred during the colony development, (ii) the generation at which these mutations occurred, (iii) the number of single and double mutant cells in the final colony and (iv) the generation at which the transient mutator subpopulation starts (for the refined model only). A minimum of 100 realizations were performed per condition.

## Results

### Selected double mutations occur in large excess during the development of wild-type yeast colonies

We measured the rate of single and double mutations, occurring during the development of wild-type yeast colonies grown without external stresses, for three different types of mutations (**Figure 1A**). We used two genetic systems to positively select for mutants carrying i) a large segmental duplication of 150 kb on chromosome XV called *D* mutants for Duplication, (Payen et al., 2008) and ii) a reciprocal translocation between chromosomes IV and X called *T* mutants for Translocation, (Gillet-Markowska et al., 2015). We also followed the appearance of loss of function mutations of *CAN1* gene on chromosome V (called *C* mutants for [Can^R^]). These types of mutation result from different molecular pathways. The formation of the large segmental duplication relies on a Pol32-dependent replication-based mechanism (24) while the translocation results from a mitotic crossover. Both events require an homologous recombination event between the 450 bp of homology shared by two truncated heteroalleles (*URA3* for the duplication and *TRP1* for the translocation, **Figure 1A**). We found a complete absence of *T* mutant, and a 30-fold decrease of the *D* rate in a *rad52Δ* mutant background, demonstrating the importance of homologous recombination for the formation of both events (**supplementary table 2**). The *C* mutants mainly corresponded to the formation of point mutations resulting from unrepaired base misincorporation during DNA replication or base modification (tautomerization, deamination, alkylation), although larger deletions or truncations could also occur. We felt that using three types of mutations based on different molecular pathways (*D*, *T* and *C*) would enable us to assess the overall level of genome instability that cells undergo during colony development.

We performed fluctuation assays on structured media to estimate mutation rates. Briefly, individual cells were plated on solid synthetic complete medium at equal distances from each other to avoid growth interference, and incubated at 30°C for 5 days, during which the cells divided for a total of 27 generations (**Figure 1B**). Subsequently, the cells of the individual colonies were plated on selective media to score the number of single and double mutants that accumulated during the initial colony development. We found that the *D*, *T* and *C* mutants occurred at rates of 3.3 x 10^-6^, 1.8 x 10^-8^ and 1.6 x 10^-7^ mutations per cell per division, respectively (**Figure 2A, supplementary table 2**). The *D* and *C* rates measured in colonies were similar to the rates obtained for cells grown in liquid medium (**supplementary table 2**) and to previously published ones (24, 28, 30).

**Figure 2:**
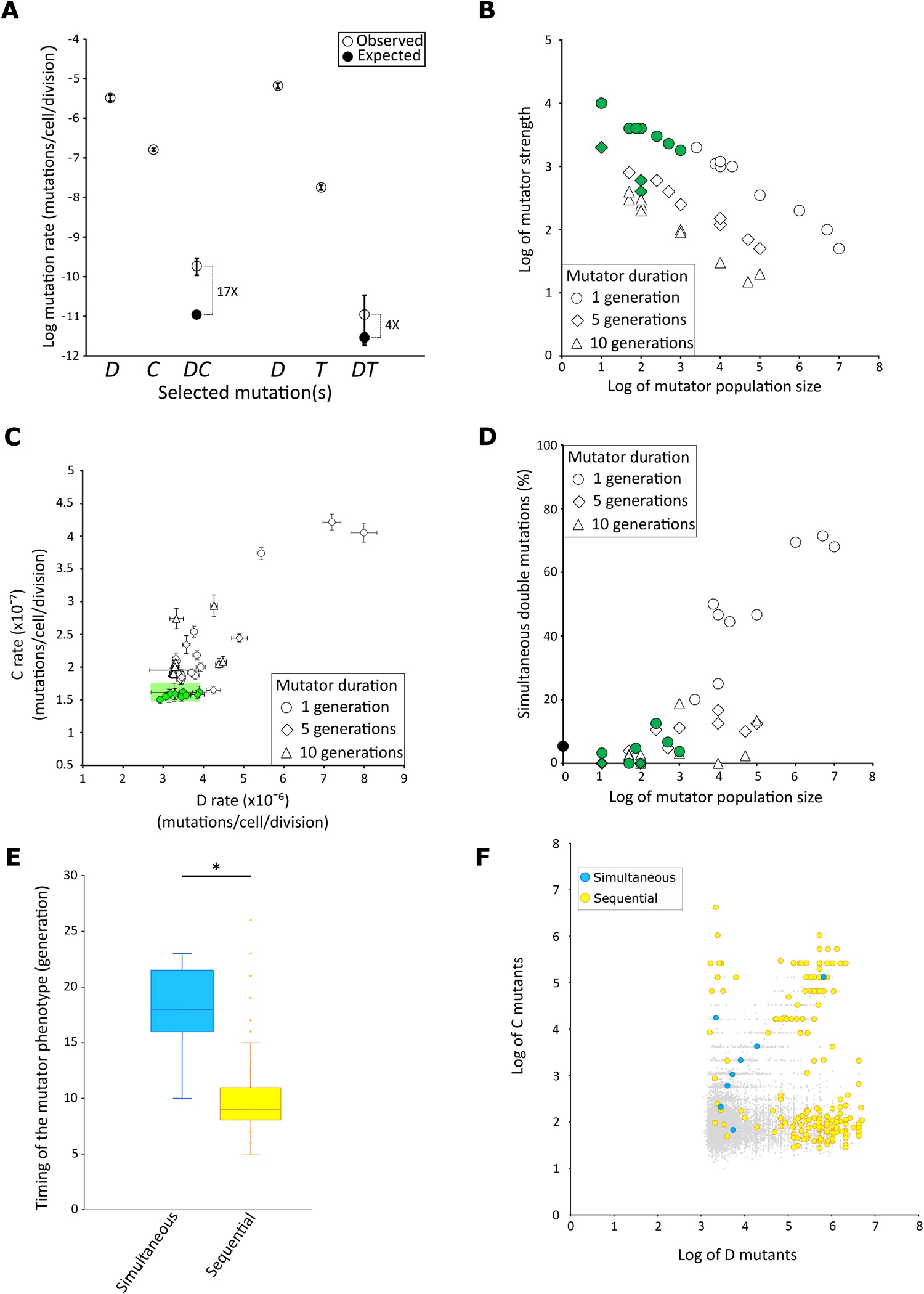
Heterogeneity of mutation rates and mutation accumulation regimes in a yeast colony. **A)** The experimentally measured single mutation rates for the segmental Duplication (*D*), Canavanine resistance (*C*) and reciprocal Translocation (*T*), and double mutation rates for Duplication and Canavanine resistance (*DC*) and Duplication and Translocation (*DT*) are symbolized by open circles. The theoretical double mutation rates (*DC*) and (*DT*) estimated with a null model of mutation accumulation are indicated by closed circles. The fold changes between the observed and theoretical double mutation rates are indicated on the side. Error bars represent the 95% likelihood ratio confidence intervals. **B)** Exploration of the parameter space of the refined model of mutation accumulation population size, mutator strength and duration of the mutator episode. The mutator subpopulation size represents the number of cells that experience the transient mutator phenotype. The mutator strength is expressed as a fold change increase over the average cell mutation rate. The three mutator episodes of 1, 5 or 10 generations are symbolized by open circles, diamonds and triangles, respectively. All reported values recapitulate the experimentally observed *DC* double mutation rate. The points colored in green correspond to combinations of size, strength and duration of the mutator phenotype that recapitulate both the observed *DC* double mutation rate and the observed single *D* and *C* mutation rates as determined by the area highlighted in green in Figure 2C. For each point at least 1,000 realizations of the refined model were performed to calculate the mutation rate. **C)** The *D* and *C* single mutation rates obtained from the realizations of the refined model for combinations of parameters of size, strength and duration of the mutator subpopulation presented in Figure 2B. Open circles, diamonds and triangles correspond to the one in Figure 2B. For each point at least 1,000 realizations were performed to calculate the mutation rates. The area highlighted in green corresponds to the 95% likelihood ratio confidence intervals of the experimentally measured *D* and *C* single mutation rates. The realizations that fall within this area recapitulate the experimental single mutation rates and correspond to the green points in Figure 2B. **D)** Percentage of simultaneously obtained *DC* mutations in the realizations of the refined model for combinations of parameters of size, strength and duration of the mutator subpopulation presented in Figure 2B. The black circle on the y-axis represents the percentage of simultaneous double mutants obtained from 55,000 realizations of the null model (*i.e.* without mutator subpopulations). The symbols colored in green are the same as in Figure 2B and C. **E)** The generation at which the mutator phenotype occurred in the realization with the refined model is represented for all realizations containing *DC* double mutants that were acquired either simultaneously (blue, n = 12) or sequentially (yellow, n = 195). * Student’s t-test p = 10^-15^. **F)** Number of single *D* and *C* mutants present in a colony after a simulated growth of 27 generations, according to the refined model ran with parameter values that recapitulate both the observed *DC* double mutation rate and the observed single *D* and *C* mutation rates, as determined in Figure 2B and C (green symbols and shaded area). Gray dots represent realizations having no double mutant after the 27 generations (n = 9,897). Blue or yellow circles represent the realizations that comprised double mutants which were acquired either simultaneously (n = 12) or sequentially (n = 195) respectively.

We showed that the presence of one type of mutation in the genome did not affect either of the other two mutation rates. Indeed, the *T* rates were not significantly different in the reference strain and in a strain that carried the segmental duplication (**supplementary figure 1A**). Likewise, the *D* rates are identical in the reference strain and in a strain carrying either the translocation or a canavanine resistant mutation. Finally, the *C* rates are also the same between the reference strain and a strain carrying the duplication (**supplementary figure 1A**). These results show that these three types of mutation are independent. We then tested whether some cells acquired the *DC* (*i.e.* coexistence in the same cell of the segmental duplication and the canavanine resistance) or the *DT* (the duplication and the translocation) double mutations during colony development by plating the cells on media selecting for the concomitant presence of two mutations. To assess this question, a large number of individual colonies were plated to estimate the double mutant appearance rates (550 and 1,006 colonies plated to select for *DC* and *DT* mutants, respectively**, supplementary table 2**). Overall, we obtained 16 *DC* and 2 *DT* mutants. As *DT* mutants were already rare, the selection for the *TC* double mutant was not attempted based on the single *T* and *C* single mutation rate (1.8 x 10^-8^ and 1.6 x 10^-7^). We checked that the growth rate of *DT* and *DC* double mutants was similar to that of the reference strain, ruling out the possibility that a fitness defect or advantage might have biased the appearance of these mutants. Note that only one double mutant *DC8* showed an increased generation time from about 120 to 180 minutes **(supplementary figure 1B**). *DC8* was not able to grow on a non-fermentable carbon source, and genome sequencing revealed the absence of mitochondrial DNA, indicating that it was a *rho0* petite mutant (**supplementary figure 1C**). Based on these numbers, the observed rates of double mutant formation was 1.8 x 10^-10^ and 1.1 x 10^-11^ for *DC* and *DT* respectively (**Figure 2A**). Because the three types of mutations are independent, the rates of double mutations occurring simultaneously during the same cell division should correspond to the product of the corresponding single mutation rates. However, we found that the observed rates of double mutant formation largely exceeded the product of the single mutation rates (340- and 90-fold higher for *DC* and *DT* rates, respectively), suggesting that the sequential acquisition of the two mutations in different cell divisions and/or an unexplained increased rate of simultaneous double mutants during the development of the colony could significantly contribute to the observed double mutation rates.

### Heterogeneous mutation rates coexist within yeast colonies

To determine expected double mutation rates that would take into account both the simultaneous occurrence of the two mutations in the same cell division, at a rate corresponding to the product of the single mutation rates, but also the sequential accumulation of the two mutations in different cell cycles during the development of the colony, we developed a null model of mutation accumulation that simulates the appearance of mutations during colony development. This model was developed with the following assumptions: i) cell growth is exponential, ii) no cell death is considered, iii) mutations appear only in daughter cells, iv) the two mutation rates are constant and identical for all cells in the colony v) reversion mutations are ignored. This model is based on only three parameters, the two single mutation rates that we experimentally measured and the number of cell generations during the development of the colony. We performed realizations of the model to estimate the expected double mutation rate and compared it with the observed double mutation rates that we experimentally determined. We found that the observed *DC* double mutation rate was 17 times higher than the expected rate (*p* = 1.1 x 10^-16^, **Figure 2A**). A similar trend was found for *DT* double mutants as the observed rate was 4 times higher (*p* = 0.04) than its expected value (**Figure 2A**). The statistical significance of this result is however low due to the small number of *DT* double mutants that we were able to isolate in the reference background (only 2 *DT* double mutants out of 1006 plated colonies), showing that the mutation rate to obtain a *DT* double mutant is at the detection limit of our experimental system.

The excess of *DC* (and to a lesser extent *DT*) double mutations is not compatible with a constant and homogenous mutation rate shared by all cells of the colony but instead reveals that heterogeneous mutation rates must coexist within the colony. We hypothesize that a subpopulation of mutator cells that generate double mutants at an increased rate would be produced during the development of a wild-type yeast colony grown in the absence of external stress.

### Transient phenotypic mutators generate double mutations

The excess of double mutants could originate either from stable genetic mutators that would have acquired mutations in genes involved in genome stability or from transient phenotypic mutators that would be genetically wild-type but express a transient mutator phenotype. We reasoned that if the double-mutants originated from genetic mutators, they would stably inherit an increased mutation rate because suppressor reversion that would restore the wild-type mutation rate should be extremely rare. However, if the double mutants originated from transient mutators they would return to a wild-type mutation rate after the 27 generations that are required to isolate them on the selective media. To determine if the double mutants came from stable or transient mutators, we measured the canavanine resistance rate in the two *DT* double mutants and found no higher mutation rate compared to the parental strain (**supplementary figure 1D**). We also measured the reciprocal translocation rate in the 16 *DC* mutants and again observed no difference in the mutation rate between the mutants and the parental strain (**supplementary figure 1D)**. These results showed that the double mutants have wild-type mutation rates, strongly suggesting that they originated from wild-type cells that expressed a transient mutator episode during the normal development of the colony.

### The subpopulation of transient mutators is small and expresses a brief yet intense mutator phenotype

In order to gain insight into possible scenarios leading to the excess of double mutant, we refined our stochastic model of mutation accumulation by implementing the formation of a subpopulation of mutator cells during the development of the colony. This mutator population was defined by its size (in number of cells), mutator strength (represented by the fold increase over the observed single mutation rates) and duration of the mutator phenotype (in number of generations). In this model, cells are randomly picked in the population to enter a mutator state in which they and their progeny stay during the entire mutator episode. We used the refined model to explore the parameters space to estimate the values that produce a double mutation rate identical to the observed *DC* double mutation rate, while keeping the single mutation rates unchanged. We set the duration of mutator phenotype to 1, 5 and 10 generations and varied the subpopulation size from 10 to 10^7^ cells and the mutator strength from 10- to 10^4^-fold increase. We plotted, for the three durations, all combinations of mutator sizes and strengths that recapitulated the experimentally determined *DC* double mutation rate (**Figure 2B**). As expected, mutator strength correlates negatively with mutator size, since a smaller subpopulation would require a higher mutator strength to achieve the observed *DC* double mutation rate. For these combinations of parameters, we then recalculated the single *D* and *C* mutation rates using the simulated data and compared them to the observed single *D* and *C* mutation rates that were experimentally measured (**Figure 2C, green shaded area**). We found that the implementation of the mutator subpopulations could modify the single mutation rates as many combinations of parameters led to a significant increase of the recalculated single mutation rates as compared to the observed single mutation rates (**Figure 2C**). As mutation rates are kept low to avoid deleterious mutation, it has been proposed that transient mutator subpopulations should not affect the average mutation rates (9). In this framework, we therefore focused on the subset of combinations of size, duration and mutator strength that did not significantly increase the recalculated single mutation rates compared to those measured experimentally. For most of these combinations, the distributions of double mutants also remained compatible with the observed ones (**Figure 2B and C green symbols, and supplementary table 3**). These corresponded to situations where the mutator durations remained short (1 or 5 generations), the mutator population sizes were small (from ten to thousands of cells) and the mutator strengths were high (from hundreds to 10^4^-fold increase, **Figure 2B, green symbols**). The same characteristics were found for the subpopulations at the origin of the *DT* double mutations (**supplementary figure 2**). Therefore, in the scenario where the mutator subpopulation should not affect the average mutation rate, the observed bursts of mutations likely result from small subpopulations of hypermutator cells.

### The majority of double mutations appear sequentially during the colony development

As double mutants can either be the product of sequential or simultaneous mutations we explored the regime of acquisition of mutations in the population by performing at least 1,000 realizations for each combination of parameters that recapitulates the observed *DC* double mutation rate. It is interesting to note that all the parameter combinations for which simultaneously acquired mutations predominate involved large mutator population sizes which resulted in single mutation rates higher than those measured experimentally, and were therefore not compatible with our observations (**Figure 2C and D**). When considering only the combinations of sizes, durations and mutator strengths that also fit the single mutation rates (**Figure 2B and C, green symbols**), simultaneously acquired mutations represented about 3%, and only up to 12.5% of all mutations, the majority of the double mutants resulting from sequentially acquired mutations (**Figure 2D**). The model also allows us to determine the timing of appearance of the mutations during the development of the colony. Firstly, for all realizations with a combination of parameters that are consistent with the observed single and double mutation rates and that possess double mutants after 27 generations, we determined at which generation the subpopulation acquired the mutator phenotype. We observed that for populations having sequentially acquired mutations, the mutator phenotype appeared rather early during the development of the colony (on average at the 9^th^ generation) while for simultaneously acquired mutation it appeared later (on average at the 17^th^ generation (**Figure 2E**). Secondly, we plotted, for each realization, the number of *D* mutants against the number of *C* mutants in the simulated colony (**Figure 2F**). After 27 generations the vast majority of the simulated colonies (98%) contained only single mutant cells (gray dots on **Figure 2F**). Additionally, 91 % of the colonies contained more *D* than *C* single mutants, in accordance with the higher mutation rate for *D* than for *C* (3.3 x 10^-6^ *vs* 1.6 x 10^-7^ mutations/cell/division, respectively). Most sequentially acquired double mutants are found in colonies encompassing a large number of *D* (> 10^5^) and/or C (> 10^4^) single mutants (**Figure 2F**). On the contrary, most of the simultaneous double mutants are found in colonies comprising a small number (< 10^4^) of both *D* and *C* single mutants (**Figure 2F**). These results suggest that the sequentially double mutant might emerge from a brief mutator episode responsible for an early acquisition of a first mutation (mainly *D* due to its higher mutation rate), and later, the second mutation would occur in the progeny of the first mutant, independently from the initial, or any, mutator episode, while, most simultaneously acquired mutations would result from later mutator episodes (**Figure 2E and F**).

### Transient mutators can generate genome-wide bursts of genomic instability

To gain more insight into the two possible regimes of mutation accumulation, *i.e.* sequential or simultaneous, we analyzed the genomes of the *DC* and *DT* double mutants, reasoning that no additional mutation should be found upon the sequential accumulation of the two mutations whereas their simultaneous appearance could be associated with a general burst of genomic instability. We karyotyped the 2 *DT* and 16 *DC* double mutants using PFGE and found, as expected, the selected reciprocal translocation between chromosomes IV and X and the large duplication on chromosome XV (**supplementary figure 3A**). We found one double mutant, the *DC3* strain, that harbored an additional band on the gel, possibly corresponding to a significantly smaller chromosome V (**supplementary figure 3A**). We also amplified and sequenced the *CAN1* gene, located on chromosome V, in the 16 *DC* mutants. For all *DC* mutants, except for *DC3* for which no PCR amplification product was obtained, the sequencing results revealed single point mutations in the *CAN1* gene, generating a frameshift or a missense mutation **(Table 1)**. These mutations are in proportion similar to those published in two large screens for canavanine resistant mutants (28, 31). We then fully sequenced, using Oxford Nanopore Technology, and *de novo* assembled the genome of 9 *DC* mutants to look for additional structural variants. We identified all the expected genetic markers present in the parental strain as well as the large segmental duplication on chromosome XV. Additionally, we identified an extra deletion of 140 bp on chromosome XIII in *DC2* and a terminal deletion of 30 kb on chromosome V in *DC3*, (**table 1 and Supplementary figure 3B)**, consistent with its PFGE karyotype (**supplementary figure 3A**). The breakpoint of the 30 kb deletion was located within the *CAN1* gene where a new telomere was added using the 5’-GGTG-3’/5’CACC-3’ sequence in *CAN1* gene as a seed for telomerase (**supplementary figure 3B**). It is noteworthy that such a case of telomere healing was never observed among the 227 *CAN1* alleles resulting from a screen for canavanine resistant mutant but is similar to the ones selected in the gross chromosomal rearrangement assay (28, 32, 33). Telomere healing at the *CAN1* locus was shown to occur at a rate of about 4 x 10^-10^ mutations per cell per division (34, 35). We used our null model with the rates of telomere healing and segmental duplication as parameters and estimated the probability to obtain 1 double mutant (*D* and terminal deletion) to be of 1.2 x 10^-11^, *i.e.* 460 lower than the rate at which we obtained the *DC3* mutant. Finally, using our long reads sequencing data we looked for point mutations in the double mutants, only keeping mutations with a quality score higher than the lowest quality score of the validated *CAN1* mutations. We found 1, 2 and 8 additional point mutations in the genomes of *DC8, DC2 and DC3* mutants, respectively (**Table 1 and supplementary table 4**). Using a point mutation rate of 3.6 × 10^−10^ substitutions per site per generation (36), we estimated that our mutants should have accumulated on average 0.2 mutation during the ∼50 generations of our experiments, *i.e.* 5, 10 and 40 times lower than what we observed for these 3 *DC* mutants. These additional substitutions are not clustered but scattered on different chromosomes, suggesting that they would result from a genome-wide mutational instability. These results suggest that at least 2 double mutants (*DC2* and *DC3*) underwent a genome-wide burst of genomic instability. They could possibly represent cases where the two selected mutations occurred simultaneously, resulting from a pathological cellular context in which mutation can accumulate in a burst in a single (or a few) cell cycle(s). Altogether, the genomic analysis of the double mutants are consistent with our model predictions which suggest that two different types of transient mutator regimes, resulting in either sequential or simultaneous double mutants, would coexist during colony development.

**Table 1:**
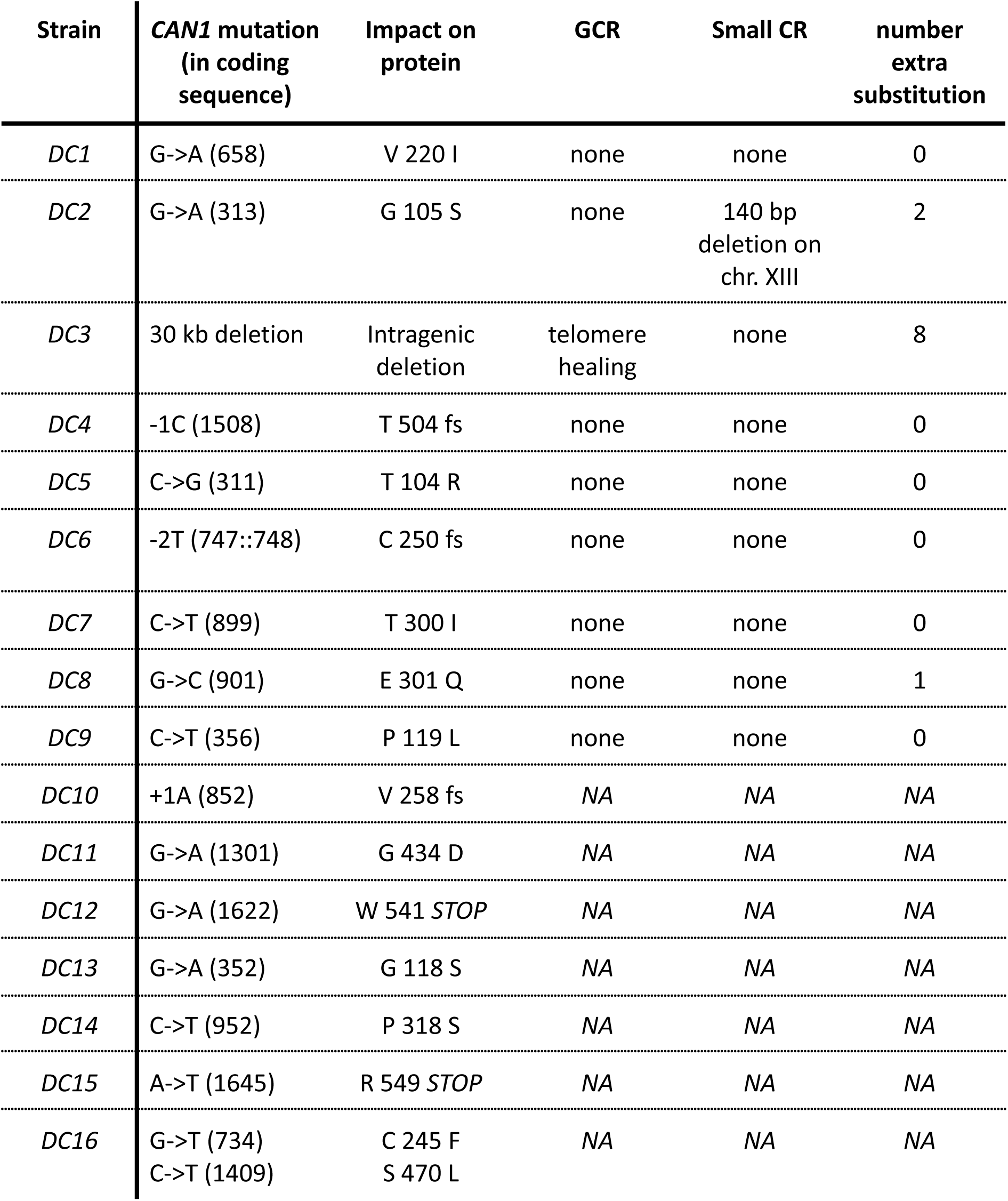
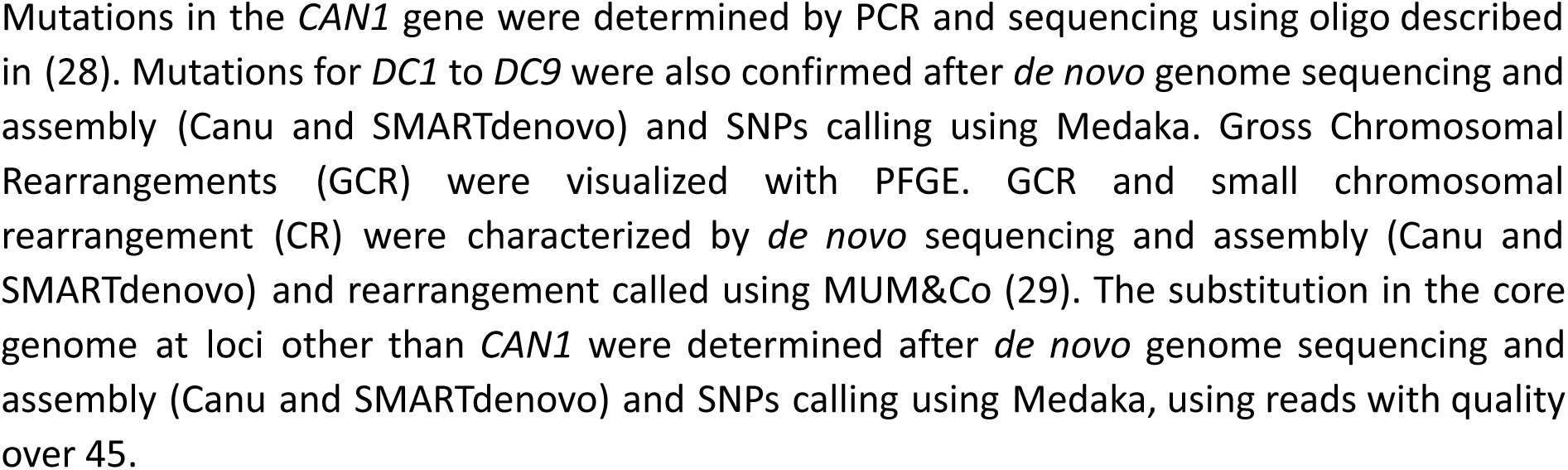
Summary of the mutation in the DC mutants.

### Transient mutator episodes also occur in replication stress conditions

As replication stress is a major contributor to genomic instability (37–39), we tested whether impaired DNA replication could be responsible for the onset of the transient mutator episodes. We measured single and double mutation rates in presence of 100 mM of hydroxyurea (HU) or in a *rad27Δ* mutant. HU is an inhibitor of the synthesis of the deoxyribonucleotides resulting in a delayed activation of replication origins and an extended S-phase (Alvino et al. 2007), while Rad27/Fen1 is involved in the processing of Okazaki fragments during DNA replication, in base excision and in mismatch repair pathways (40, 41). We found that HU stimulates the formation of all single and double mutational events with *D, C* and *DC* rates being 2, 3 and 12 times higher than in the control, respectively (**Figure 3 and supplementary table 2**). Moreover, the observed rate of double mutants was 25 times higher than expected, as compared to the 17 times increase in the absence of HU (**Figure 3**). Similarly, in a *rad27Δ* mutant, both single *D* and *C* and the double *DC* mutation rates were increased 2, 20 and 200 times, respectively (**figure 3 and supplementary table 2**) and the observed rate of double mutants was 27 times higher than expected (**Figure 3**). Thus, while mutation rates showed an increase under the influence of potent replication stressors like HU treatment and a *rad27Δ* mutation, they remained significantly lower than the strength exhibited by the transient mutators that we previously estimated to be in the order of 4 x 10^2^ to 10^4^-fold increase. Additionally, the elevated ratio of observed to expected double mutant formation rate in both HU-treated and *rad27Δ* cells is similar to the one observed in the reference and suggests a cumulative effect of the replication stress and the unknown causative determinants responsible for the transient mutator subpopulation. We used the model to estimate the potential size and strength of the transient mutator subpopulation in HU-treated and *rad27Δ* cells. We set the transient mutator duration for 1 or 5 generations and selected parameter combinations that recapitulated the observed single and double mutation rates in these conditions (**supplementary figure 4**). The results showed the size and strength of the transient mutator subpopulations were in the same range as in the reference strain, suggesting that adding the replication stressors did not affect the transient mutator subpopulation (**supplementary figure 4**). All these results showed that the transient mutator subpopulation would remain mostly unchanged in colonies grown under a systemic replication stress, thus suggesting that other determinants would be at the origin of the transient mutator subpopulations.

**Figure 3:**
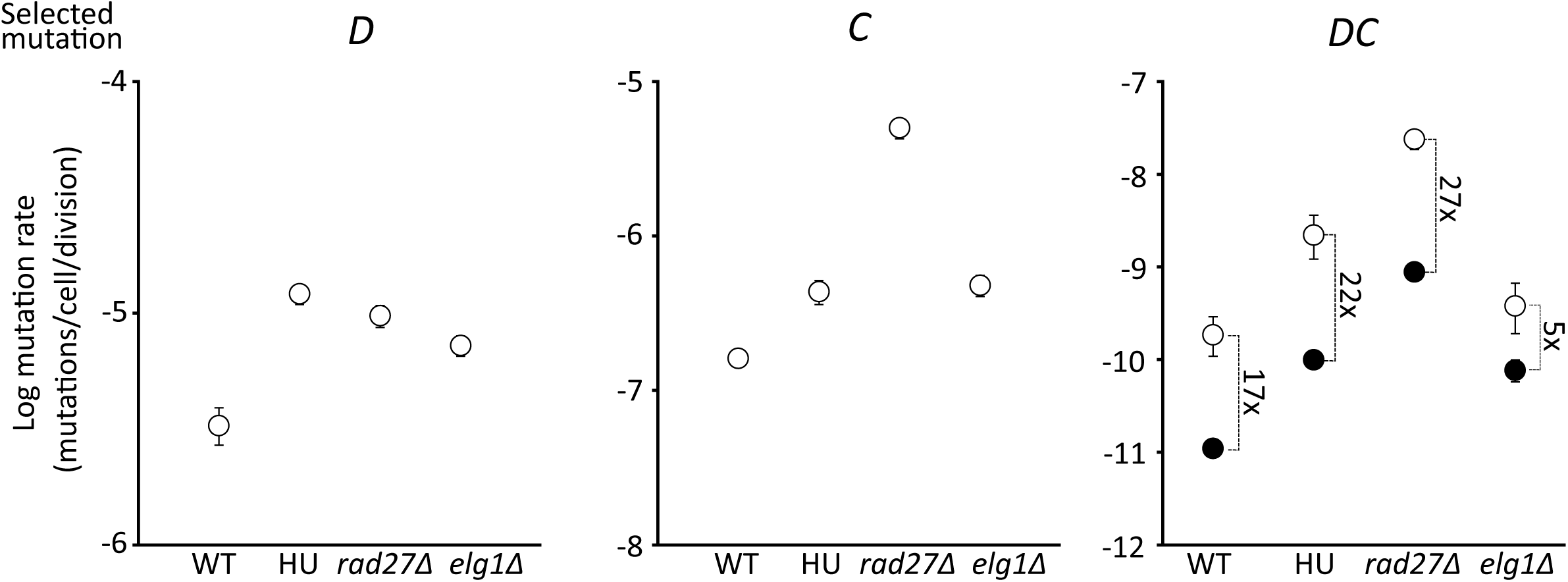
Single and double mutation rates in replication stress conditions. The experimentally measured single *D* and *C* and double *DC* mutation rates are symbolized by open circles. The theoretical double *DC* mutation rates, estimated from the null model of mutation accumulation after 27 generations, are indicated by closed circles. The fold changes between the measured and theoretical double mutation rates are indicated on the side. Error bars represent the 95% likelihood ratio confidence intervals. The genetic backgrounds of the strains and the presence of HU (100 mM) are indicated below the x-axis.

### The ELG1-dependent recombination salvage pathway contributes to the double mutation excess

Elg1/ATAD5 is a key protein at the interface of the replication and repair processes. It is part of the Elg1-RLC complex whose function is to unload SUMOylated forms of the processivity replication factor PCNA. In an *elg1* mutant background, the accumulation of SUMOylated PCNA on the chromatin generates a global replication defect, genome instability and increased mutagenesis (42–45). Additionally, Elg1 activity is required for an alternative DNA damage repair mechanism called “salvage recombination” (Arbel et al. mBio 2020). We measured the *D*, *T* and *C* single mutation rates in an *elg1*Δ mutant background and found, as expected, increased rates as compared to the reference strain (**Figure 3 and supplementary table 2**). These rates were close to those measured in the presence of 100 mM HU, thus, the expected *elg1*Δ *DC* double mutation rate predicted by the stochastic model was also similar to the one in the presence of HU (**Figure 3**). However, we found a 5 times lower observed *DC* double mutation in the *elg1*Δ background as compared to the one in HU-treated cells (**Figure 3 and supplementary table 2**). Additionally, the *elg1*Δ observed *DC* double mutation rate was only 5 times higher than the expected rate, thus lower than the 17-fold increase found in the wild-type strain (**Figure 3**). Using the stochastic model, we determined the parameters that would reproduce the observed *DC* double mutation rates in an *elg1*Δ mutant. The results showed that for a mutator state lasting 1 or 5 generations the observed *DC* mutation rate in an *elg1*Δ mutant can be produced by a similar mutator subpopulation size as in the reference strain but with a lower mutation rate (**supplementary figure 4**). The size of the transient mutator subpopulation being equal, the *DC* mutants in excess in wild-type compared to *elg1Δ* might result from the recombination salvage pathway at stalled replication forks (46).

## Discussion

This study aimed to characterize the mutation accumulation regime in isogenic cell populations. Our results revealed the presence of transient mutator subpopulations during the development of yeast colonies grown in complete medium in absence of external stresses, reinforcing the possibility that heterogeneity of mutation rate in isogenic populations could be the norm rather than the exception (Alexander et al., 2017; Matic, 2019). Our data indicate that these transient mutator subpopulations would be small and undergo short yet very intense mutational bursts, resulting in rate increases ranging from several hundreds to several thousands folds (**Figure 2**). These mutator subpopulations could be at the limit of cellular life sustainability given that constitutively high mutation rates decrease the fitness of cells and can lead to lineage extinction due to highly deleterious mutations in essential genes (8, 12, 47). It is possible that some combinations of simulation parameters matching our observed data, particularly those with the highest mutation rates, are not consistent with cell survival. However, in good agreement with the duration and strength of the mutator episodes that we reported here, it was shown that haploid yeast cells could withstand over a thousand fold increase of their mutation rate for several generations before experiencing visible growth defects due to mutation accumulation (Herr et al., 2011).

Our findings are based on the assumption that the contribution of the transient mutators to the single mutations would be small, implying that the single mutation rates recalculated from the realizations of the refined model should not significantly differ from the single mutation rates that we experimentally measured and used as parameter values to feed the model. However, we cannot rule out the possibility that larger mutator subpopulations, which may be less mutagenic, could be responsible for the observed excess of double mutants. However, this would imply that experimentally measured single mutation rates would include a significant contribution from single mutants produced by large subpopulations of transient mutators and that, as a result, baseline single mutation rates would actually be lower than experimentally measured rates. This hypothesis is unlikely, as several studies have suggested that single mutation rates should not be affected by the presence of transient mutator subpopulations (9, 17). Similarly, we cannot exclude a more complex scenario where several successive mutational bursts happen during colony development. Further exploration of these possibilities seems necessary for future research.

We characterized two different regimes of acquisition of double mutations and found that the main regime would be sequential (**Figure 2D**). In this regime, a precocious and transient mutator episode allows the generation of a first mutation early in the development of the colony, leading to the formation of a large subpopulation of the first mutant by clonal expansion (**Figure 2E and F**). The second mutation is acquired later, within the clonal mutant subpopulation but without the need for any mutator phenotype. This scenario is in agreement with the genome analysis of the *DC* double mutants that we recovered given that 7 out of the 9 clones did not carry any other mutation than the two selected ones (**Table 1**). We note that in human, the early acquisition of a mutation as a result of precocious mutator episodes can have huge impact in the adaptive trajectory of disease and in particular in the formation of cancer cell populations, creating mosaicism of subclones during tumorigenesis with the formation of driver and passenger mutations (1, 48, 49). The second regime corresponds to the simultaneous acquisition of the two mutations and could result from an episode of SGI. According to our stochastic model, this would occur only in a minority of cases (from 0 to 12.5% depending on parameter combinations, **Figure 2D**). In accordance with these estimates, only two out of the 9 *DC* clones (*DC2* and *DC3*) carried extra mutations, including a small internal deletion, a terminal deletion associated with a new telomere addition and multiple point mutations, spread on several chromosomes, suggesting that the mutator cells endured an SGI episode (**Table 1, Supplementary figure 3 and supplementary table 4**). SGI was previously reported in diploid yeast cells growing without external stresses, by characterizing an excess of loss of heterozygosity (LOH) events as compared to expected rates (19, 20). Thirty percent of the clones obtained after selecting for the two LOH events also carried extra chromosomal rearrangements, and it was proposed that 15% of the clones experiencing a LOH events in *S. cerevisiae* could do so by SGI (20). Interestingly, traces of SGI can potentially be found at the population level in *S. cerevisiae*. The generation of a Reference Assembly Panel, consisting of telomere-to-telomere genome assemblies for 142 strains representatives of the entire genetic diversity of the species, revealed some natural isolates with highly rearranged genomes and cases of complex aneuploidies, *i.e.* aneuploid chromosomes carrying additional large-scale genomic rearrangements such as translocations or large deletions (50). Similarly in human, episodes of SGI, allowing the acquisition of several mutations simultaneously, can also contribute to the transformation of human cells into a malignant state and account for the massive genome reorganization observed in tumor cells that could not be explained by a gradual accumulation (51–55). DC2 and DC3 clones were isolated as single unique colonies on the selective plates, indicating that the double mutations at their origin occurred at the last cell division during the colony development, suggesting that SGI resulted from late mutator episodes.

The existence of an early and a late regime of acquisition of double mutations suggests that different stresses during the development of the colony could be at the origin of the sequentially and simultaneously acquired mutations. Indeed, yeast colony formation are biphasic with first an exponential growth for about 24 generations (about 10^7^ cells) with vigorous metabolic activity as suggested by the presence of a high concentration of rDNA in the cell, followed by a second phase characterized by the expression of stress-induced genes such as *SSA3* and *HSP26* associated with a state of stress after 5 days (56, 57). The stresses that would trigger the transient mutator episodes during yeast colony development are not known but may include the stochastic partition of protein involved in DNA metabolism, transcription or translation errors generating loss of function or toxic gain of function proteins involved in DNA repair or a metabolic heterogeneity in the local environment of the cells in the colony (6, 9, 16–18). For example, the noise in gene expression of *RAD27* or *RAD52* can lead to heterogeneity in recombination rate in an isogenic population (58). Furthermore, the complex reprogramming of the genetic networks and the formation of differentiated cell populations during the colony development lead to the formation of metabolically active and stress-resistant cells as well as starving and stress-sensitive cells (59–66). The mutator subpopulations could also arise during the transition from one metabolism to another. Indeed, a spike of reactive oxygen species (ROS) has been observed during the change of carbon source or the passage to anoxia (67, 68). The molecular mechanisms by which both sequential and simultaneous double mutations are increased are not known but the deficit of double mutants in a *elg1*Δ background, compared to the reference, suggests that the ELG1-dependent recombination may play a role, possibly by counteracting the anti-recombination function of the *Srs2* helicase (**Figure 3**)(21, 69, 70).

In conclusion, our study provides a precise phenomenological dissection of the spontaneous appearance of transient mutator subpopulations during the development of wild-type yeast colonies grown in the absence of external stress. These findings have far-reaching implications in terms of genome dynamics and evolution. Further work will be needed to characterize the molecular triggers at the origin of the transient mutator bursts.

## Acknowledgements

We thank Bernard Dujon, Marina Elez, Ivan Matic and Zhou Xu for their valuable feedback on the paper. This work was supported by the Agence Nationale de la Recherche ANR-16-CE12-0019, ANR-18-CE12-0004 and ANR-20-CE12-0020.

## Supplementary Figure legends

**Supplementary figure 1.**
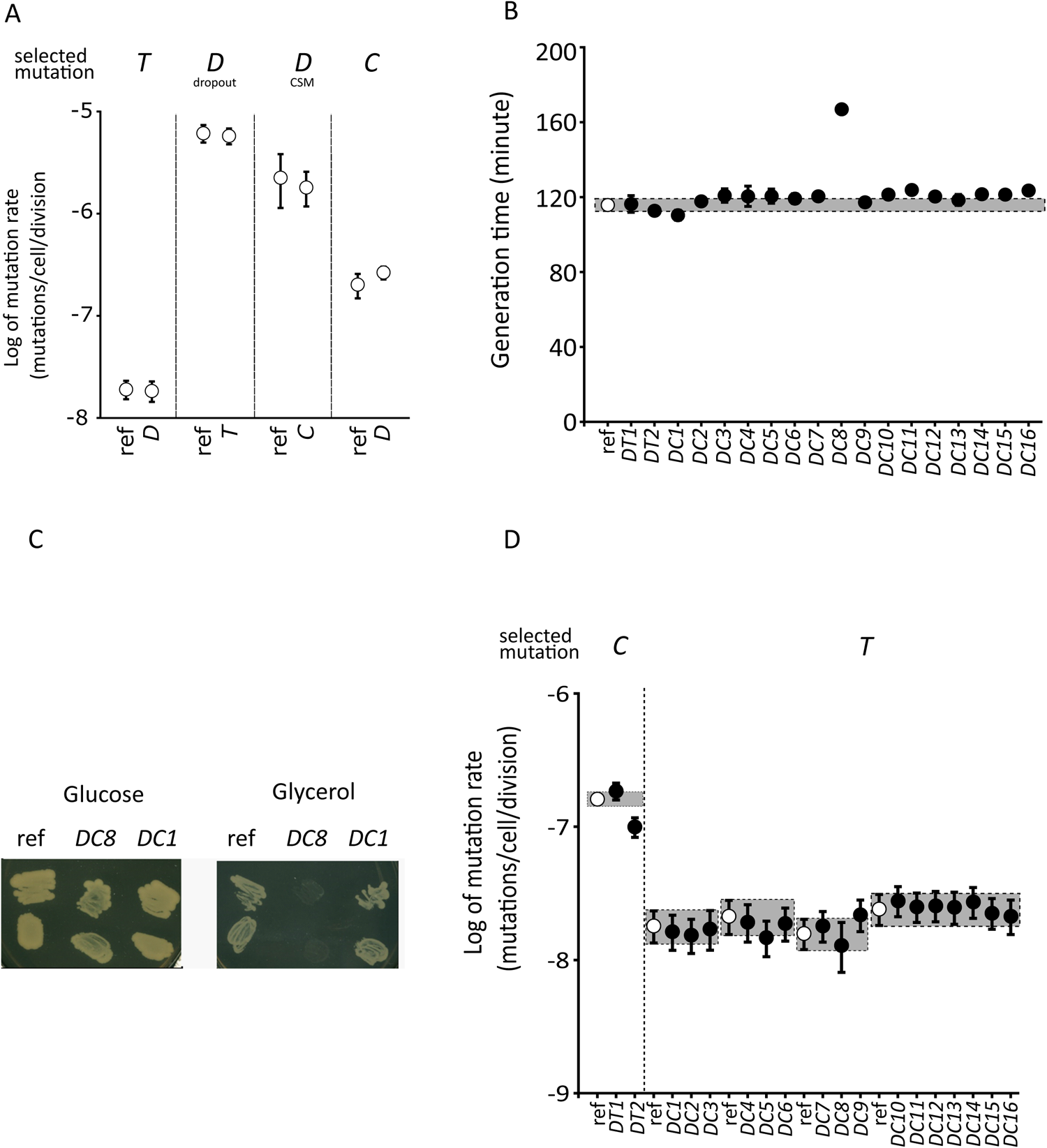
Properties of the single and double mutants. A) Comparison of mutation rates between the YAG142 reference strain (ref) and the single mutants carrying the Translocation(*T*), the segmental Duplication (*D*) or the resistance to Canavanine mutation (*C*). The single *T* rate was measured in a *D* mutant. The single *D* rate was measured in either the *T* or *C* mutants using two synthetic media, the “dropout” (Sigma-Aldrich) or the CSM (MP Biotech). The single *C* rate was measured in a *D* mutant. Error bars represent the 95% likelihood ratio confidence intervals. B) The generation time in minute of the YAG142 reference strain (ref - open circle) and the Duplication and Translocation (*DT*) or Duplication and Canavanine resistant (*DC*) double mutants (black circle), grown in YPD broth at 28°C without agitation. Each point represents the average of a minimum of three replicates. Error bars represent standard deviation to the mean. C) Cellular patches of the parental strain YAG142 (ref), *DC8* and *DC1* double mutants grown at 30°C on YP-glucose (a fermentable carbon source) and YP-glycerol (a non-fermentable carbon source). D) Comparison of mutation rates between the YAG142 reference strain (ref, open circles) and the double mutants carrying the Duplication and Translocation (*DT*) or Duplication and Canavanine resistant (*DC*, black circles). The single *C* and *T* rates were measured in the 2 *DT* and 16 *DC* double mutants, respectively. The *DC s*trains were tested in four different batches with each time the reference strain used as control. Error bars represent the 95% likelihood ratio confidence intervals.

**Supplementary figure 2:**
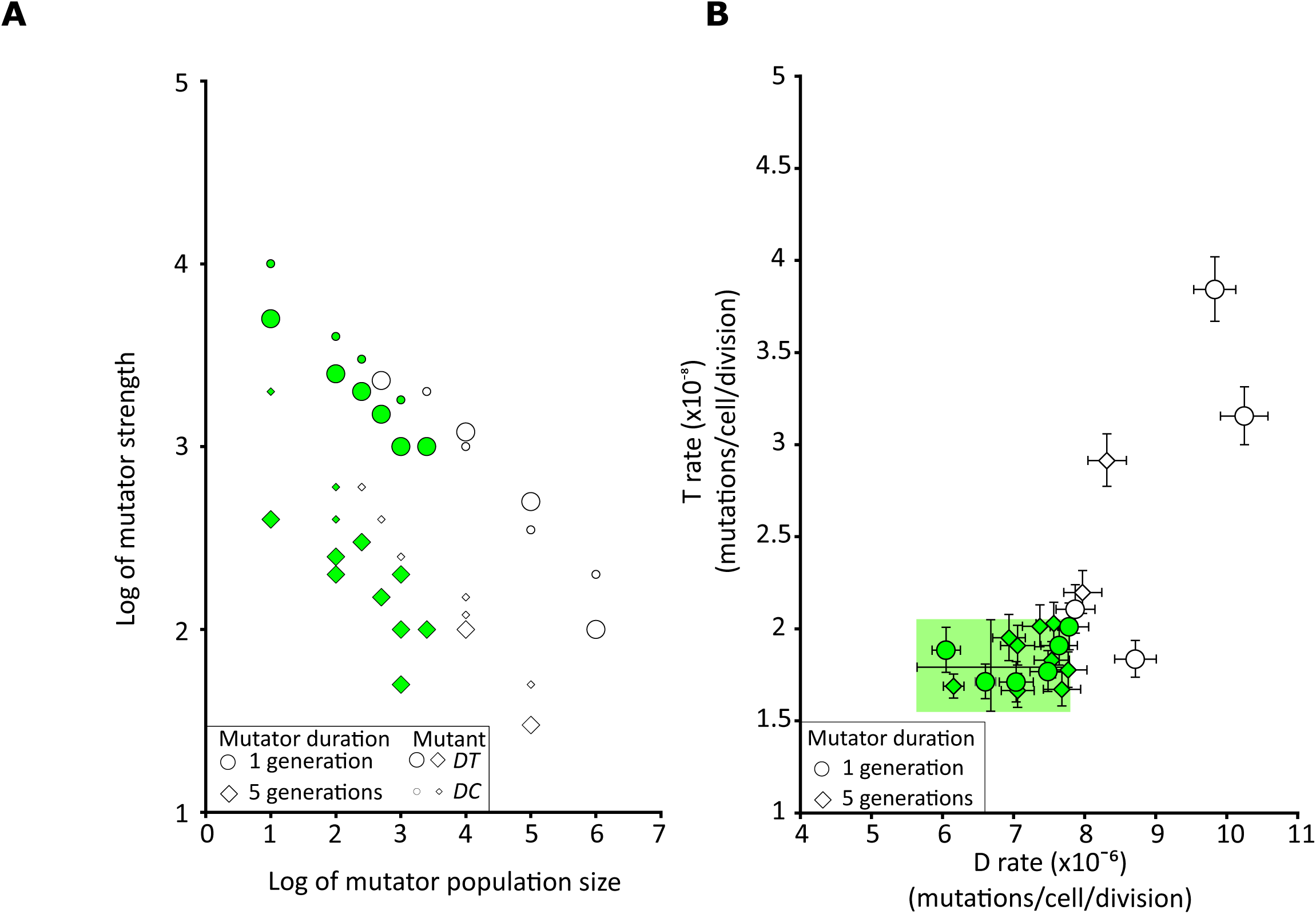
Exploration of parameter space for the duration, size and strength of the mutator subpopulation for the formation of DT double mutants. **A)** The mutator subpopulation size represents the number of cells that experience the transient mutator phenotype. The mutator strength is expressed as a fold change increase of the general cell mutation rate. The two mutator durations of 1 and 5 generations are symbolized by open circles and diamonds, respectively. The symbol size refers to *DT* (big) and *DC* (small) double mutants. All reported values recapitulate the experimentally observed *DT* and *DC* double mutation rates. For each point at least 1,000 realizations of the refined model were performed to calculate the mutation rate. The points colored in green correspond to combinations of size, strength and duration of the mutator phenotype that recapitulate both the observed *DT* and *DC* double mutation rates and the observed single *D*, *T* and *C* mutation rates as determined by the area highlighted in green in the B panel. **B)** The symbols are as in the A panel. For each point at least 1,000 realizations have been performed to calculate the mutation rates. The area highlighted in green corresponds to the 95% likelihood ratio confidence intervals of the experimentally measured *D* and *T* single mutation rates. The realizations that fall within this area recapitulate the experimental single mutation rates and therefore were colored in green in the A panel.

**Supplementary figure 3:**
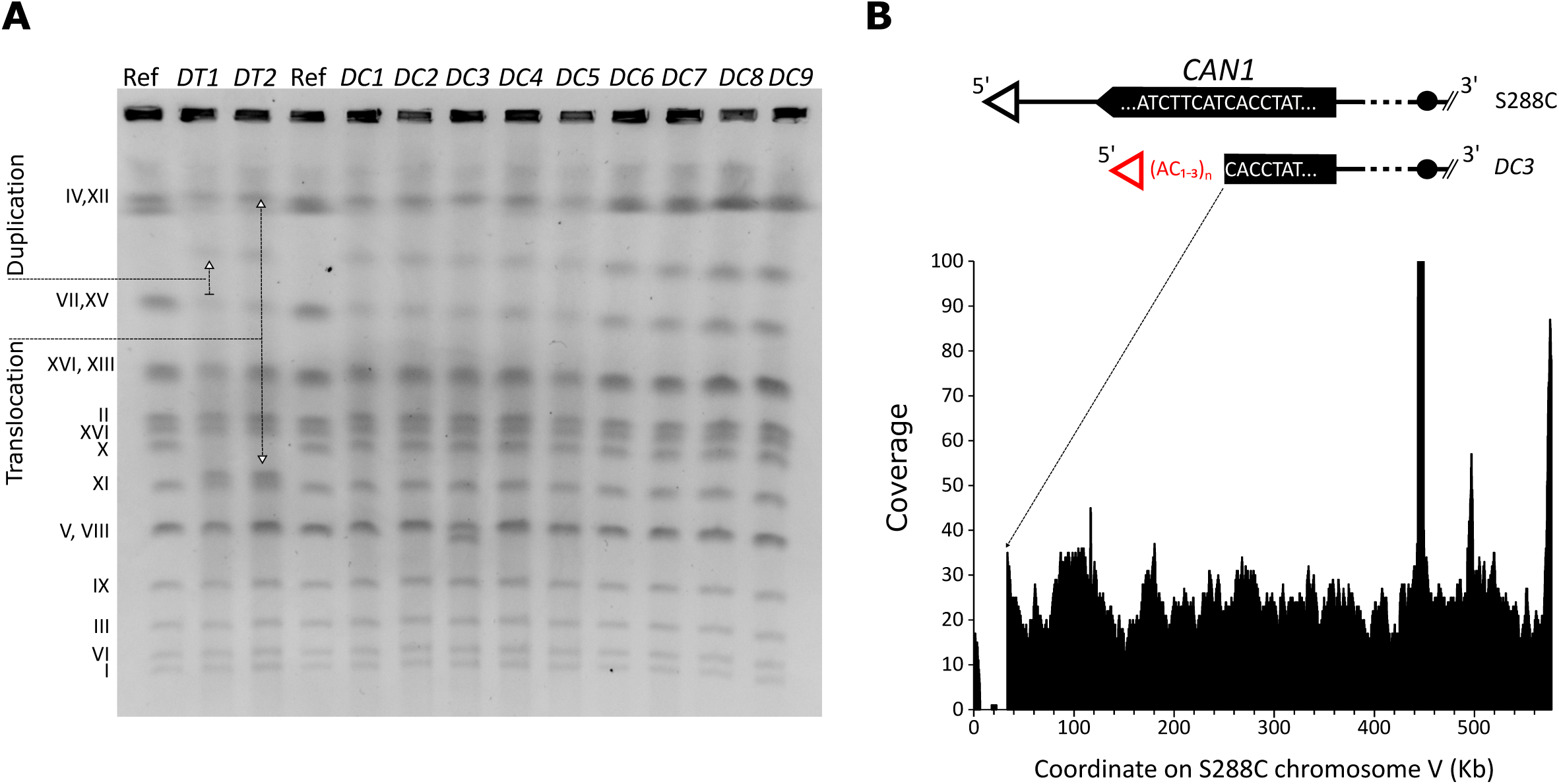
Characterization of an additional large deletion in the DC3 double mutant. **A)** Pulsed-field gel electrophoresis of the parental strain YAG142 (Ref), 2 *DT* and 9 *DC* double mutants. The bands characterizing the selected reciprocal translocation and segmental duplication are indicated by arrows. The *DC3* karyotype shows a smaller band corresponding to either chromosome V or VIII. **B)** The DNA coverage obtained from *de novo* genome sequencing along the chromosome V of the *DC3* strain. Upper part shows a schematic of chromosome V left arm in the reference strain S288C and in the *DC3* double mutant at the *CAN1* locus. The deletion occurred within the *CAN1* gene and a new telomere was added using a 5’-GGTG-3’/CACC-3’ seed.

**Supplementary figure 4:**
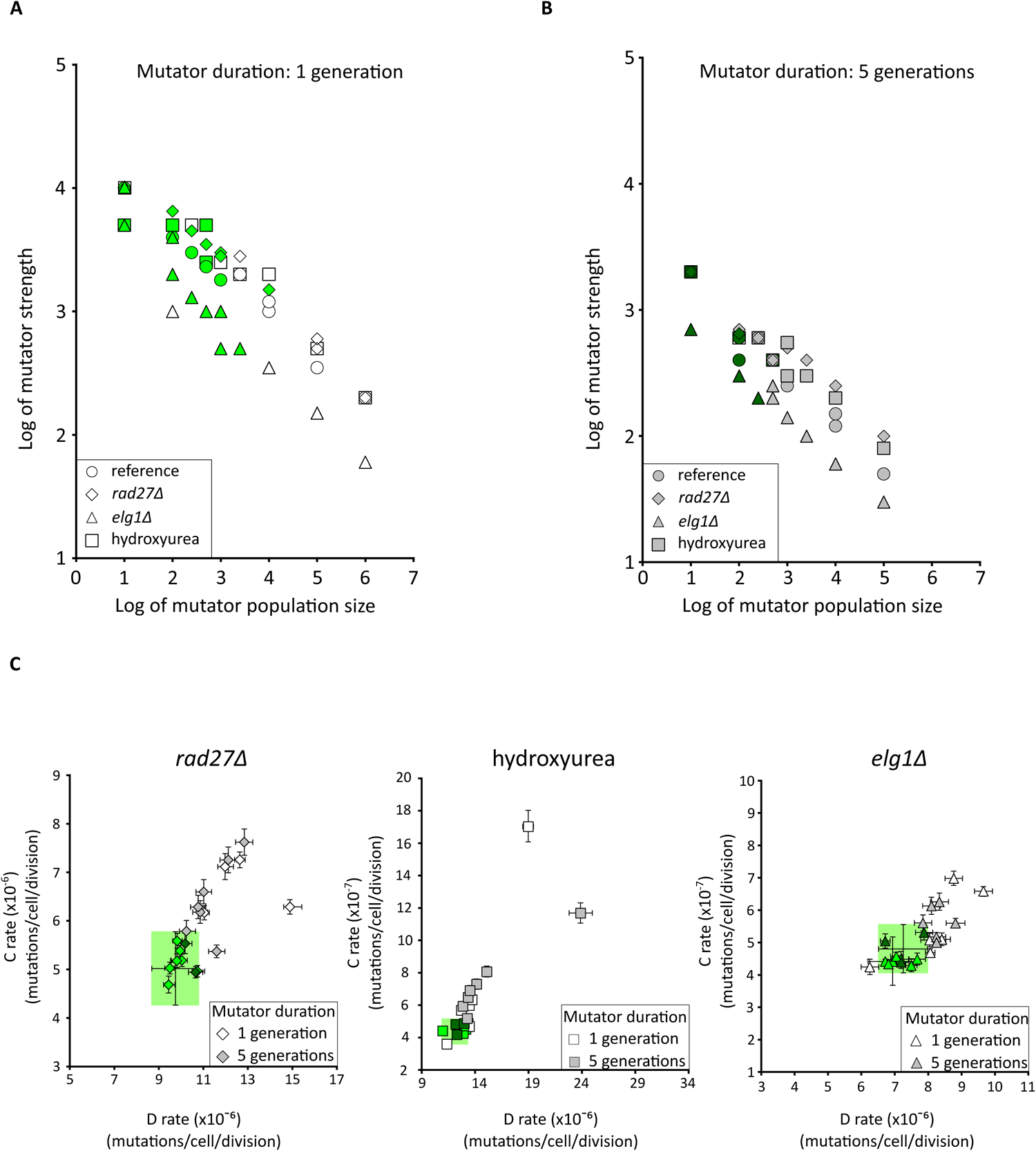
Mutator population size and strength in HU treated cells, rad27Δ and elg1Δ mutants. **A)** Combination of parameters of mutator size and strength for a mutator episode of one generation. The mutator subpopulation size represents the number of cells that experience the transient mutator phenotype. The mutator strength is expressed as a fold change increase of the general cell mutation rate. All reported values recapitulate the experimentally observed *DC* double mutation rate. The points colored in green correspond to combinations of size and strength of the mutator phenotype that recapitulate both the observed *DC* double mutation rates and the single *D* and *C* mutation rates as determined by the area highlighted in green in the C panel. For each point at least 1,000 realizations of the refined model were performed to calculate the mutation rate **B)** Combination of parameters of mutator size and strength for a mutator episode of five generations. The symbols are identical to those of panel A. **C)** The symbols are as in the A and B panels. For each point at least 1,000 realizations were performed to calculate the mutation rates. The area highlighted in green corresponds to the 95% likelihood ratio confidence intervals of the experimentally measured *D* and *C* single mutation rates. The realizations that fall within this area recapitulate the experimental single mutation rates and therefore were colored in green in the A panel.

## Supplementary Tables

**Supplementary table 1:**
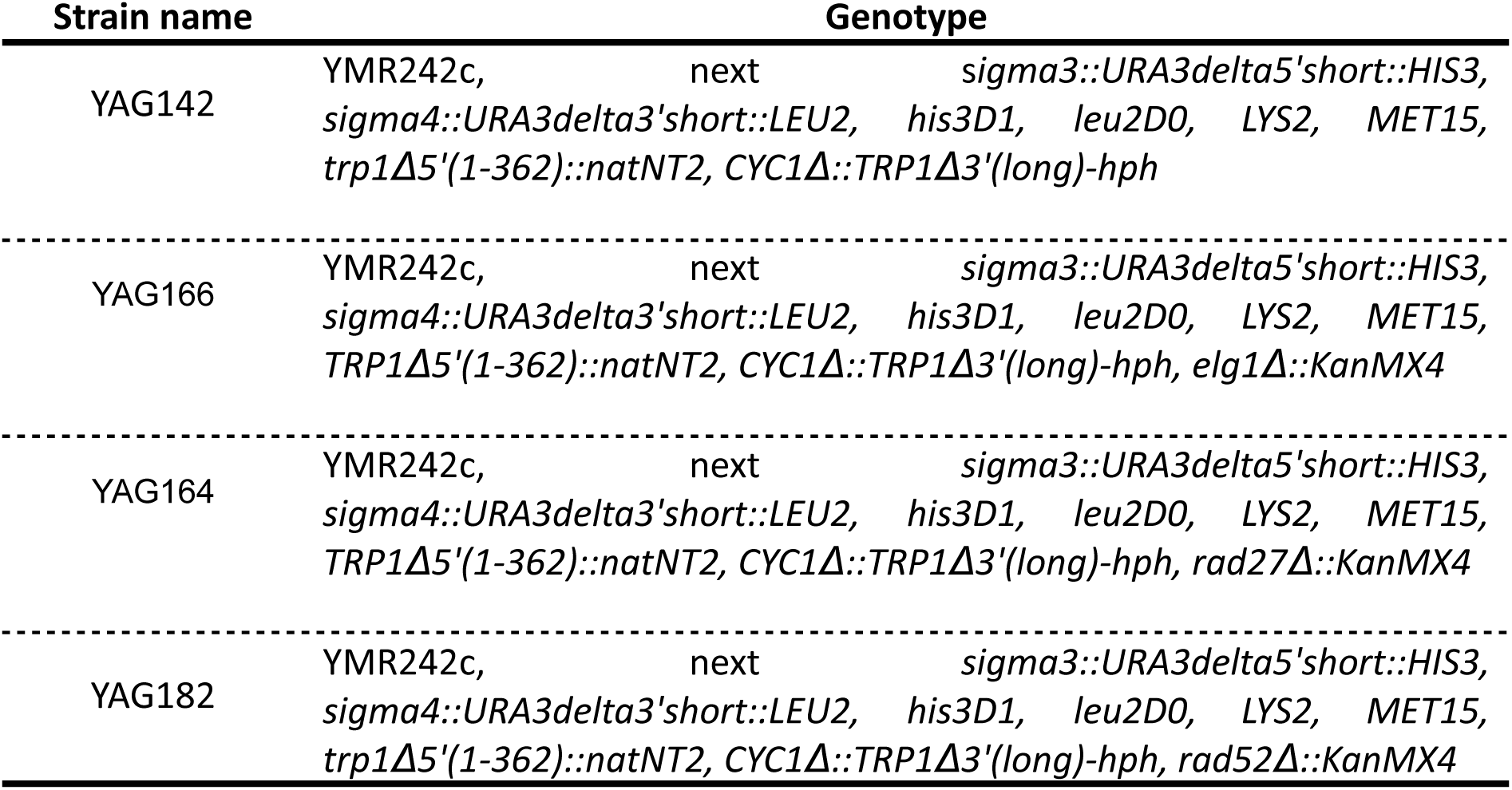
Strains table.

**Supplementary table 2:**
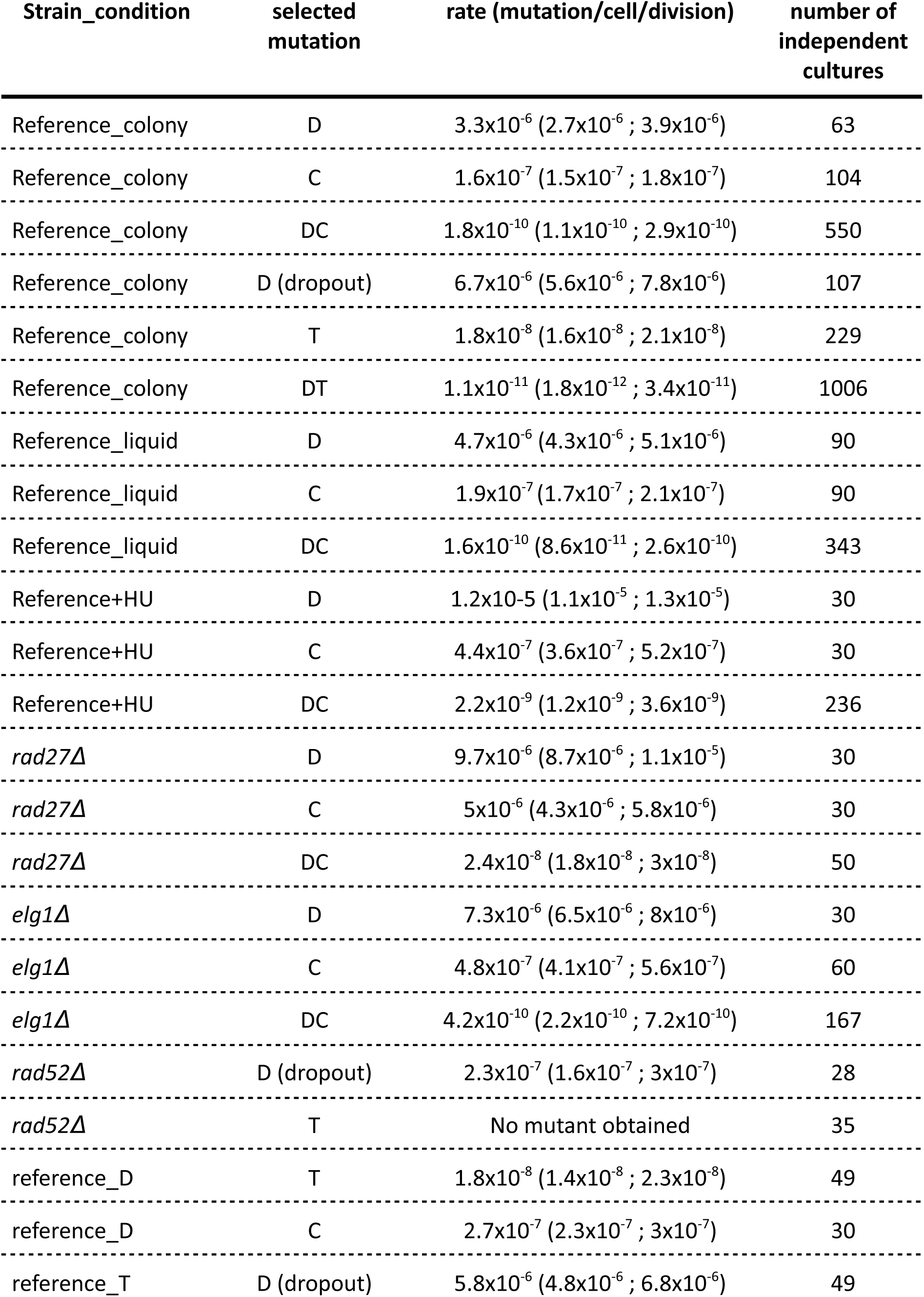
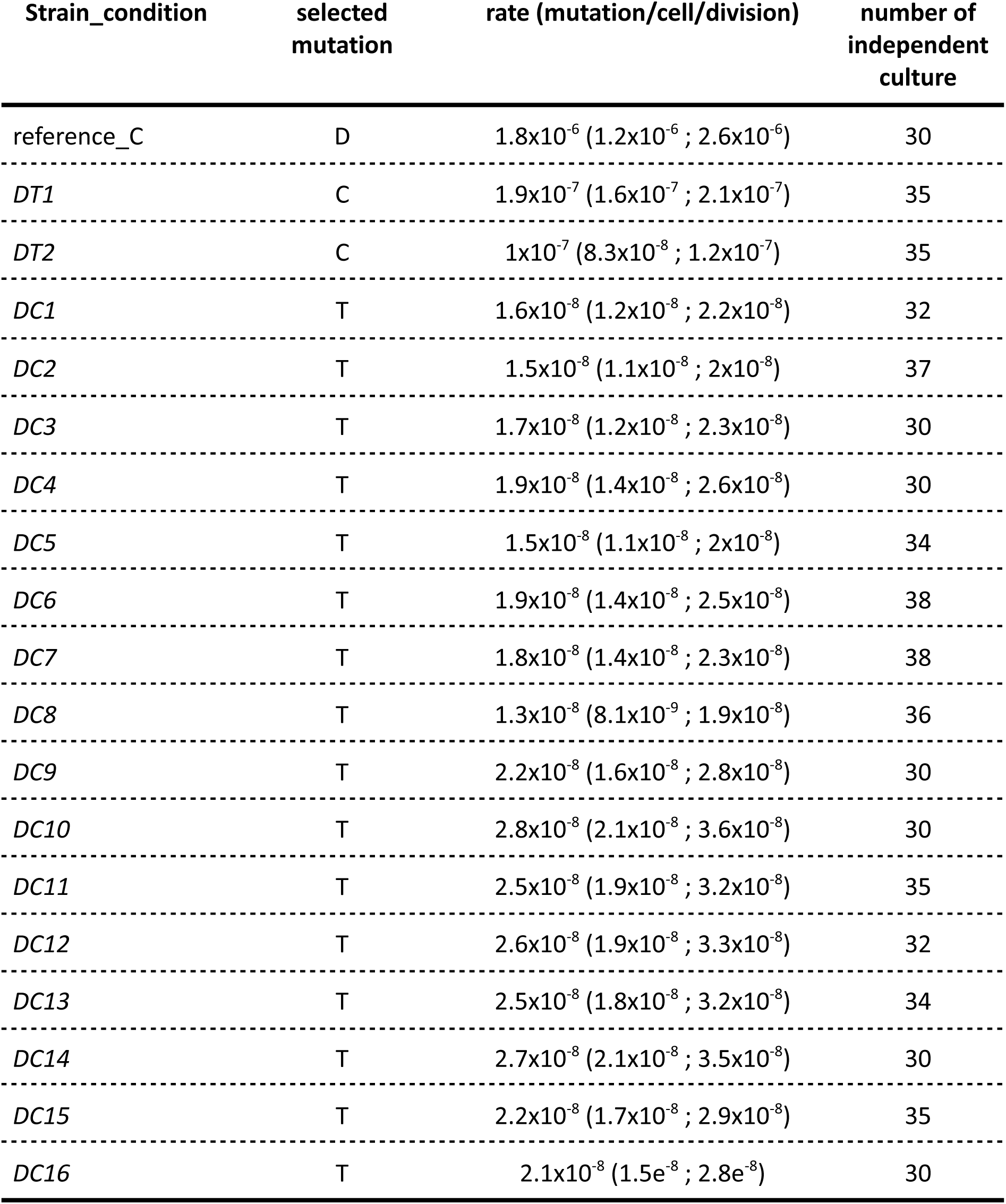
Single and double mutation rate measured for this study.

**Supplementary table 3:** Comparison of the distribution of double mutants obtained between the simulation and experimentally observed.

| Mutator Duration<br>(in generation) | Mutator population<br>size | Mutator increase<br>fold change | *p*<br>obs vs simu |
| --- | --- | --- | --- |
| 1 | 10 | 10000 | 0.08 |
| 1 | 50 | 4000 | 0.99 |
| 1 | 75 | 4000 | 0.28 |
| 1 | 100 | 4000 | 0.7 |
| 1 | 250 | 3000 | 0.04 |
| 1 | 500 | 2300 | 0.91 |
| 1 | 1000 | 1800 | 0.7 |
| 5 | 10 | 2000 | 0.04 |
| 5 | 100 | 400 | 0.47 |
| 5 | 100 | 600 | 0.7 |

**Supplementary table 4:** Non selected substitution mutations in DC2, DC3 and DC8.

| Strain | Mutation | #chromosome (position) |
| --- | --- | --- |
| DC2 | C->T | chr. II (543996) |
| DC2 | A->G | chr. IX (196591) |
| DC3 | C->T | chr. IV (317835) |
| DC3 | A->T | chr. VII (94265) |
| DC3 | C->T | chr. VIII (27951) |
| DC3 | C->T | chr. VIII (381360) |
| DC3 | A->G | chr. XIII (764215) |
| DC3 | G->T | chr. XIV (42862) |
| DC3 | G->A | chr. XIV (499069) |
| DC3 | T->A | chr. XVI (630865) |
| DC8 | G->A | chr. V (556341) |

The presence of the substitutions in the core genome were determined after *de novo* genome sequencing and assembly (Canu and SMARTdenovo) and SNPs calling using Medaka, using reads with quality over 45. The positions are the one given by the assembly performed with Canu.

## Notes

### Competing Interest Statement

The authors have declared no competing interest.

### Summary of Updates

mostly text changes

## References

1. L. A. Loeb, Human cancers express mutator phenotypes: origin, consequences and targeting. Nat. Rev. Cancer 11, 450–457 (2011).

2. H. Liu, J. Zhang, The rate and molecular spectrum of mutation are selectively maintained in yeast. Nat. Commun. 12, 4044 (2021).

3. M. Lynch, et al., Genetic drift, selection and the evolution of the mutation rate. Nat. Rev. Genet. 17, 704–714 (2016).

4. P. D. Sniegowski, P. J. Gerrish, T. Johnson, A. Shaver, The evolution of mutation rates: separating causes from consequences. BioEssays 22, 1057–1066 (2000).

5. L. R. Heasley, N. M. V. Sampaio, J. L. Argueso, Systemic and rapid restructuring of the genome: a new perspective on punctuated equilibrium. Curr. Genet. 67, 57–63 (2021).

6. J. W. Drake, Too Many Mutants with Multiple Mutations. Crit. Rev. Biochem. Mol. Biol. 42, 247–258 (2007).

7. H. K. Alexander, S. I. Mayer, S. Bonhoeffer, Population Heterogeneity in Mutation Rate Increases the Frequency of Higher-Order Mutants and Reduces Long-Term Mutational Load. Mol. Biol. Evol. 34, 419–436 (2017).

8. A. Giraud, The rise and fall of mutator bacteria. Curr. Opin. Microbiol. 4, 582–585 (2001).

9. I. Matic, Mutation Rate Heterogeneity Increases Odds of Survival in Unpredictable Environments. Mol. Cell 75, 421–425 (2019).

9. T. Swings, et al., Adaptive tuning of mutation rates allows fast response to lethal stress in Escherichia coli. eLife 6, e22939 (2017).

10. E. Denamur, I. Matic, Evolution of mutation rates in bacteria. Mol. Microbiol. 60, 820–827 (2006).

11. A. J. Herr, et al., Mutator Suppression and Escape from Replication Error–Induced Extinction in Yeast. PLOS Genet. 7, e1002282 (2011).

12. C. Zeyl, M. Mizesko, J. A. de Visser, Mutational meltdown in laboratory yeast populations. Evol. Int. J. Org. Evol. 55, 909–917 (2001).

13. J. P. Pribis, et al., Gamblers: An Antibiotic-Induced Evolvable Cell Subpopulation Differentiated by Reactive-Oxygen-Induced General Stress Response. Mol. Cell 74, 785–800.e7 (2019).

14. D. van Dijk, et al., Slow-growing cells within isogenic populations have increased RNA polymerase error rates and DNA damage. Nat. Commun. 6, 7972 (2015).

15. A. C. Woo, L. Faure, T. Dapa, I. Matic, Heterogeneity of spontaneous DNA replication errors in single isogenic Escherichia coli cells. Sci. Adv. 4, eaat1608 (2018).

16. J. Ninio, Transient mutators: a semiquantitative analysis of the influence of translation and transcription errors on mutation rates. Genetics 129, 957–962 (1991).

17. S. Uphoff, et al., Stochastic activation of a DNA damage response causes cell-to-cell mutation rate variation. Science 351, 1094–1097 (2016).

18. L. R. Heasley, R. A. Watson, J. L. Argueso, Punctuated Aneuploidization of the Budding Yeast Genome. Genetics 216, 43–50 (2020).

19. N. M. V. Sampaio, et al., Characterization of systemic genomic instability in budding yeast. Proc. Natl. Acad. Sci. U. S. A. 117, 28221–28231 (2020).

20. M. Arbel, A. Bronstein, S. Sau, B. Liefshitz, M. Kupiec, Access to PCNA by Srs2 and Elg1 Controls the Choice between Alternative Repair Pathways in Saccharomyces cerevisiae. mBio 11, e00705–20 (2020).

21. C. Baker Brachmann, et al., Designer deletion strains derived from Saccharomyces cerevisiae S288C: A useful set of strains and plasmids for PCR-mediated gene disruption and other applications. Yeast 14, 115–132 (1998).

22. A. Wach, A. Brachat, R. Pöhlmann, P. Philippsen, New heterologous modules for classical or PCR-based gene disruptions in Saccharomyces cerevisiae. Yeast 10, 1793–1808 (1994).

23. C. Payen, R. Koszul, B. Dujon, G. Fischer, Segmental Duplications Arise from Pol32-Dependent Repair of Broken Forks through Two Alternative Replication-Based Mechanisms. PLOS Genet. 4, e1000175 (2008).

24. A. Gillet-Markowska, G. Louvel, G. Fischer, bz-rates: A Web Tool to Estimate Mutation Rates from Fluctuation Analysis. G3 GenesGenomesGenetics 5, 2323–2327 (2015).

25. Q. Zheng, rSalvador: An R Package for the Fluctuation Experiment. G3 GenesGenomesGenetics 7, 3849–3856 (2017).

26. T. Török, D. Rockhold, A. D. King, Use of electrophoretic karyotyping and DNA-DNA hybridization in yeast identification. Int. J. Food Microbiol. 19, 63–80 (1993).

27. G. I. Lang, A. W. Murray, Estimating the per-base-pair mutation rate in the yeast Saccharomyces cerevisiae. Genetics 178, 67–82 (2008).

28. S. O’Donnell, G. Fischer, MUM&Co: accurate detection of all SV types through whole-genome alignment. Bioinformatics 36, 3242–3243 (2020).

29. M.-E. Huang, A.-G. Rio, A. Nicolas, R. D. Kolodner, A genomewide screen in Saccharomyces cerevisiae for genes that suppress the accumulation of mutations. Proc. Natl. Acad. Sci. 100, 11529–11534 (2003).

30. P. Jiang, et al., A modified fluctuation assay reveals a natural mutator phenotype that drives mutation spectrum variation within Saccharomyces cerevisiae. eLife (2021) 10.7554/eLife.68285 (October 6, 2022).

31. V. Pennaneach, C. D. Putnam, R. D. Kolodner, Chromosome healing by de novo telomere addition in Saccharomyces cerevisiae. Mol. Microbiol. 59, 1357–1368 (2006).

32. C. D. Putnam, V. Pennaneach, R. D. Kolodner, Chromosome healing through terminal deletions generated by de novo telomere additions in Saccharomyces cerevisiae. Proc. Natl. Acad. Sci. 101, 13262–13267 (2004).

33. C. Chen, R. D. Kolodner, Gross chromosomal rearrangements in Saccharomyces cerevisiae replication and recombination defective mutants. Nat. Genet. 23, 81–85 (1999).

34. A. Piazza, et al., Stimulation of Gross Chromosomal Rearrangements by the Human CEB1 and CEB25 Minisatellites in Saccharomyces cerevisiae Depends on G-Quadruplexes or Cdc13. PLOS Genet. 8, e1003033 (2012).

35. A. Serero, C. Jubin, S. Loeillet, P. Legoix-Né, A. G. Nicolas, Mutational landscape of yeast mutator strains. Proc. Natl. Acad. Sci. 111, 1897–1902 (2014).

36. R. D. Kolodner, C. D. Putnam, K. Myung, Maintenance of Genome Stability in Saccharomyces cerevisiae. Science 297, 552–557 (2002).

37. L. Toledo, K. J. Neelsen, J. Lukas, Replication Catastrophe: When a Checkpoint Fails because of Exhaustion. Mol. Cell 66, 735–749 (2017).

38. M. K. Zeman, K. A. Cimprich, Causes and consequences of replication stress. Nat. Cell Biol. 16, 2–9 (2014).

39. F. A. Calil, et al., Rad27 and Exo1 function in different excision pathways for mismatch repair in Saccharomyces cerevisiae. Nat. Commun. 12, 5568 (2021).

40. D. X. Tishkoff, N. Filosi, G. M. Gaida, R. D. Kolodner, A Novel Mutation Avoidance Mechanism Dependent on S. cerevisiae RAD27 Is Distinct from DNA Mismatch Repair. Cell 88, 253–263 (1997).

41. M. Bellaoui, et al., Elg1 forms an alternative RFC complex important for DNA replication and genome integrity. EMBO J. 22, 4304–4313 (2003).

42. T. Kubota, K. Myung, A. D. Donaldson, Is PCNA unloading the central function of the Elg1/ATAD5 replication factor C-like complex? Cell Cycle 12, 2570–2579 (2013).

43. K. Lee, S. H. Park, Eukaryotic clamp loaders and unloaders in the maintenance of genome stability. Exp. Mol. Med. 52, 1948–1958 (2020).

44. K. Shemesh, et al., A structure–function analysis of the yeast Elg1 protein reveals the importance of PCNA unloading in genome stability maintenance. Nucleic Acids Res. 45, 3189–3203 (2017).

45. C. P. Lehmann, A. Jiménez-Martín, D. Branzei, J. A. Tercero, Prevention of unwanted recombination at damaged replication forks. Curr. Genet. 66, 1045–1051 (2020).

46. P. Funchain, et al., The Consequences of Growth of a Mutator Strain of Escherichia coli as Measured by Loss of Function Among Multiple Gene Targets and Loss of Fitness. Genetics 154, 959–970 (2000).

47. M. Mohiuddin, R. F. Kooy, C. E. Pearson, De novo mutations, genetic mosaicism and human disease. Front. Genet. 13 (2022).

48. Z. Seferbekova, A. Lomakin, L. R. Yates, M. Gerstung, Spatial biology of cancer evolution. Nat. Rev. Genet. 24, 295–313 (2023).

49. S. O’Donnell, et al., Telomere-to-telomere assemblies of 142 strains characterize the genome structural landscape in Saccharomyces cerevisiae. Nat. Genet. 55, 1390–1399 (2023).

50. S. C. Baca, et al., Punctuated Evolution of Prostate Cancer Genomes. Cell 153, 666–677 (2013).

51. P. J. Campbell, et al., Pan-cancer analysis of whole genomes. Nature 578, 82–93 (2020).

52. Y. Li, et al., Patterns of somatic structural variation in human cancer genomes. Nature 578, 112–121 (2020).

53. F. Pellestor, Chromoanagenesis: cataclysms behind complex chromosomal rearrangements. Mol. Cytogenet. 12, 6 (2019).

54. E. Rheinbay, et al., Analyses of non-coding somatic drivers in 2,658 cancer whole genomes. Nature 578, 102–111 (2020).

55. J.-R. Meunier, M. Choder, Saccharomyces cerevisiae colony growth and ageing: biphasic growth accompanied by changes in gene expression. Yeast 15, 1159–1169 (1999).

56. L. Váchová, M. Čáp, Z. Palková, Yeast Colonies: A Model for Studies of Aging, Environmental Adaptation, and Longevity. Oxid. Med. Cell. Longev. 2012, e601836 (2012).

57. J. Liu, J.-M. François, J.-P. Capp, Gene Expression Noise Produces Cell-to-Cell Heterogeneity in Eukaryotic Homologous Recombination Rate. Front. Genet. 10 (2019).

58. M. Čáp, L. Štěpánek, K. Harant, L. Váchová, Z. Palková, Cell Differentiation within a Yeast Colony: Metabolic and Regulatory Parallels with a Tumor-Affected Organism. Mol. Cell 46, 436–448 (2012).

59. M. Čáp, L. Váchová, Z. Palková, Yeast Colony Survival Depends on Metabolic Adaptation and Cell Differentiation Rather Than on Stress Defense*. J. Biol. Chem. 284, 32572–32581 (2009).

60. C. Correia-Melo, et al., Cell-cell metabolite exchange creates a pro-survival metabolic environment that extends lifespan. Cell 186, 63–79.e21 (2023).

61. S. Kamrad, et al., Metabolic heterogeneity and cross-feeding within isogenic yeast populations captured by DILAC. Nat. Microbiol. 8, 441–454 (2023).

62. N. Paczia, et al., Extensive exometabolome analysis reveals extended overflow metabolism in various microorganisms. Microb. Cell Factories 11, 122 (2012).

63. Z. Palková, et al., Ammonia Pulses and Metabolic Oscillations Guide Yeast Colony Development. Mol. Biol. Cell 13, 3901–3914 (2002).

64. Z. Palková, L. Váchová, Spatially structured yeast communities: Understanding structure formation and regulation with omics tools. Comput. Struct. Biotechnol. J. 19, 5613–5621 (2021).

65. L. Váchová, L. Hatáková, M. Čáp, M. Pokorná, Z. Palková, Rapidly Developing Yeast Microcolonies Differentiate in a Similar Way to Aging Giant Colonies. Oxid. Med. Cell. Longev. 2013, 102485 (2013).

66. R. Dirmeier, et al., Exposure of Yeast Cells to Anoxia Induces Transient Oxidative Stress. J. Biol. Chem. 277, 34773–34784 (2002).

67. M. G. Koerkamp, et al., Dissection of Transient Oxidative Stress Response in Saccharomyces cerevisiae by Using DNA Microarrays. Mol. Biol. Cell 13, 2783–2794 (2002).

68. H. Ogiwara, A. Ui, T. Enomoto, M. Seki, Role of Elg1 protein in double strand break repair. Nucleic Acids Res. 35, 353–362 (2007).

69. S. Tamang, et al., The PCNA unloader Elg1 promotes recombination at collapsed replication forks in fission yeast. eLife 8, e47277 (2019).

